# Glycosaminoglycans are specific endosomal receptors for *Yersinia pseudotuberculosis* Cytotoxic Necrotizing Factor

**DOI:** 10.1101/2020.05.18.101790

**Authors:** Stefanie Kowarschik, Julian Schöllkopf, Thomas Müller, Songhai Tian, Julian Knerr, Hans Bakker, Stephan Rein, Min Dong, Stefan Weber, Robert Grosse, Gudula Schmidt

## Abstract

The Cytotoxic Necrotizing Factor Y (CNFY) is produced by the gram-negative, enteric pathogen *Yersinia pseudotuberculosis*. The bacterial toxin belongs to a family of deamidases, which constitutively activate Rho GTPases, thereby balancing inflammatory processes. We identified heparan sulfate proteoglycans as essential host cell factors for intoxication with CNFY. Using flow cytometry, microscopy, knockout cell lines, pulsed electron–electron double resonance and bio-layer interferometry, we studied the role of glucosaminoglycans in the intoxication process of CNFY. To analyze toxin-glucosaminoglycan interaction we utilized a truncated CNFY (CNFY_709-1014_). Especially this C-terminal part of CNFY, which encompasses the catalytic activity, binds with high affinity to heparan sulfates. CNFY binding with the N-terminal domain to its protein receptor seems to induce a first conformational change supporting the interaction between the C-terminal domain and heparan sulfates, which seems sterically hindered in the full toxin. A second conformational change occurs by acidification of the endosome, probably allowing insertion of the hydrophobic regions of the toxin into the endosomal membrane. Our findings suggest that heparan sulfates play a major role for intoxication within the endosome, rather than being relevant for an interaction at the cell surface. Lastly, cleavage of heparin sulfate chains by heparanase is likely required for efficient uptake of the toxic enzyme into the cytosol of mammalian cells.

**Author Summary:** The RhoA deamidating Cytotoxic Necrotizing Factor Y (CNFY) from *Yersinia pseudotuberculosis* is a crucial virulence factor that is important for successful infection of mammalian cells by the pathogen. The mode of action by which CNFY is able to intoxicate cells can be divided into the following steps: Binding to the cell surface, internalization, translocation from the endosome to the cytosol and deamidation of RhoA. We show, that CNFY uses heparan sulfates to maximize the amount of molecules entering the cytosol. While not being necessary for toxin binding and uptake, the sugars hold a key role in the intoxication process. We show that CNFY undergoes a conformational change at a low endosomal pH, allowing the C-terminal domain to be released from the endosomal membrane by the action of heparanase. This study reveals new insights into the CNFY-host interaction and promotes understanding of the complex intoxication process of bacterial toxins.

## Introduction

*Yersinia pseudotuberculosis* is a gram-negative, enteric pathogen, which causes self-limiting gastrointestinal infections. The bacterium produces several virulence factors like adhesins and a type-III secretion system, which injects diverse effector proteins (Yersinia outer proteins, Yops) into mammalian cells. Moreover, *Y. pseudotuberculosis* secretes the protein toxin Cytotoxic Necrotizing Factor (CNFY) via outer membrane vesicles into the environment (1). The toxin belongs to a family of deamidases modifying small GTPases of the Rho family (2, 3). Rho GTPases are molecular switches, which govern a wide variety of signaling pathways, including the rearrangement of the actin cytoskeleton, gene synthesis and survival (4). Small GTPases are regulated by binding and hydrolyzing GTP. The CNFY-catalyzed modification blocks intrinsic as well as the GAP-stimulated GTP hydrolysis, because it converts a specific glutamine (Gln63 in RhoA) required for GTP hydrolysis to glutamate, leading to constitutive activation of the targeted small GTPases (5, 6). CNFY improves translocation of Yops into host cells thereby supporting inflammatory responses of the host (7). Following binding to mammalian cells, the toxin is taken up by receptor-mediated endocytosis, however, a specific cellular toxin receptor is still not known. Subsequently, the catalytic part of the toxin is released from acidified endosomes into the cytosol where it deamidates RhoA (8).

In previous work we showed that the *Escherichia coli* toxin CNF1 (61% identity to CNFY) binds with high affinity to the Lutheran/Basal Cell Adhesion Molecule (Lu/BCAM). This interaction is required for intoxication of mammalian cells (9). In contrast, CNFY does not bind to Lu/BCAM. Previous results of cell culture experiments with pharmacological inhibition of heparin sulfation led to the assumption that proteoglycans may be involved in CNFY action (10). Here, we show that glycosaminoglycans (GAG) are involved in recruiting CNFY to mammalian cells. GAG are linear, unbranched negatively charged poly-sugars, for example heparin, heparan sulfate, chondroitin sulfate A or dermatan sulfate, covalently attached to the core protein of a proteoglycan (PG). The GAG synthesis is very complex and involves many different proteins.

The first enzyme transferring Xylose to a specific serine within the recognition sequences of the core protein is the Xylose transferase (XYLT) (11). Addition of Xylose to the core protein is required for the biosynthesis of the growing GAG chain. Heparan sulfates are synthesized by five different glucosyltransferases (EXT1, EXT2, EXTL1, EXTL2 and EXTL3) and may contain more than 100 sugar units (12). Some of the sugar units are frequently modified further for example by sulfation or epimerization, leading to generation of a wide variety of different GAG. PG are components of the extracellular matrix, the plasma membrane and are part of secretory vesicles. Main plasma membrane PG are either syndecans, which are integral membrane proteins or glypicans, attached to the surface by a glycosyl-phosphatidylinositol (GPI) anchor. Cell membrane anchored PG are well known receptors or co-receptors for a variety of macromolecules and viruses (13). Recently, sulfated glucosaminoglycans have been identified to interact with the bacterial toxin *Clostridium difficile* toxin A and to mediate its uptake into target cells (14). The aim of this study was to understand the role of PG and their GAG side chains in the CNFY intoxication process. Using flow cytometry and microscopy, mutant cell lines and bio-layer interferometry we show that heparan sulfates are crucial for CNFY-mediated intoxication of mammalian cells. They do not only serve as surface receptors, but are functionally essential components for proper toxin translocation within the endosomal compartment.

## Results

*Yersinia pseudotuberculosis* CNFY is an important virulence factor, which is crucial for successful infection of mammals. An infection with a bacterial pathogen results in strong induction of the host immune response. CNFY has been shown to influence inflammatory processes in mice (7). Therefore, we intended to analyze the toxins direct activity on immune cells.

### Immune cells are not the preferred targets of CNFY

In a first set of experiments, we studied toxin binding to different human and mouse cells. Therefore, we labeled GST-CNFY with the fluorescent marker DyLight488. Cells were incubated with the labeled toxin and washed. Bound fluorescence was then analyzed by flow cytometry. Surprisingly, tested mouse immune cells: monocytes/macrophages (RAW 264.7) and B lymphocytes (J558L) and most of the human immune cells: T lymphocytes (Molt-4) as well as B lymphocytes (Ramos) did not bind or only marginally bound the toxin (Fig. 1A). In contrast, mouse fibroblasts (MEF cells), human epithelial cells (HeLa cells) as well as human monocytes/macrophages (THP-1 cells) bound the labeled toxin. We further analyzed uptake of CNFY by directly visualizing the toxin-catalyzed modification of RhoA. Deamidated RhoA shifts to apparent higher molecular weight in urea SDS-PAGE and therefore can be distinguished from the unmodified GTPase. As expected, and in line with the results for toxin binding, CNFY was taken up into mouse MEF, human THP-1 and HeLa cells, whereas there was no shift of RhoA detectable in the other tested immune cells (Fig. S1). Our data are summarized in table 1.

**Table 1:** Immune cells are not the preferred target of CNFY. The table shows the data generated by FACS and shift assays for binding and intoxication of indicated cell lines with CNFY. Toxin binding was studied using several lines of immune cells and DL488-labeled GST-CNFY.

|  |  | HSPG<br>[mean bound<br>fluorescence] | CNFY<br>[mean bound<br>fluorescence] | RhoA<br>deamidation |
| --- | --- | --- | --- | --- |
| Murine cell lines | Monocytes/macrophages<br>(RAW 264.7) | 8 | 3 | - |
|  | B lymphocytes<br>(J558L) | 0 | 0 | - |
|  | Fibroblasts<br>(MEF) | 329 | 30 | + |
| Human cell lines | Monocytes/macrophages<br>(THP-1) | 4 | 31 | + |
|  | T lymphocytes<br>(Molt-4) | 4 | 4 | - |
|  | B lymphocytes<br>(Ramos) | 14 | 1 | - |
|  | Epithelial cells<br>(HeLa) | 56 | 45 | + |

**Figure 1:**
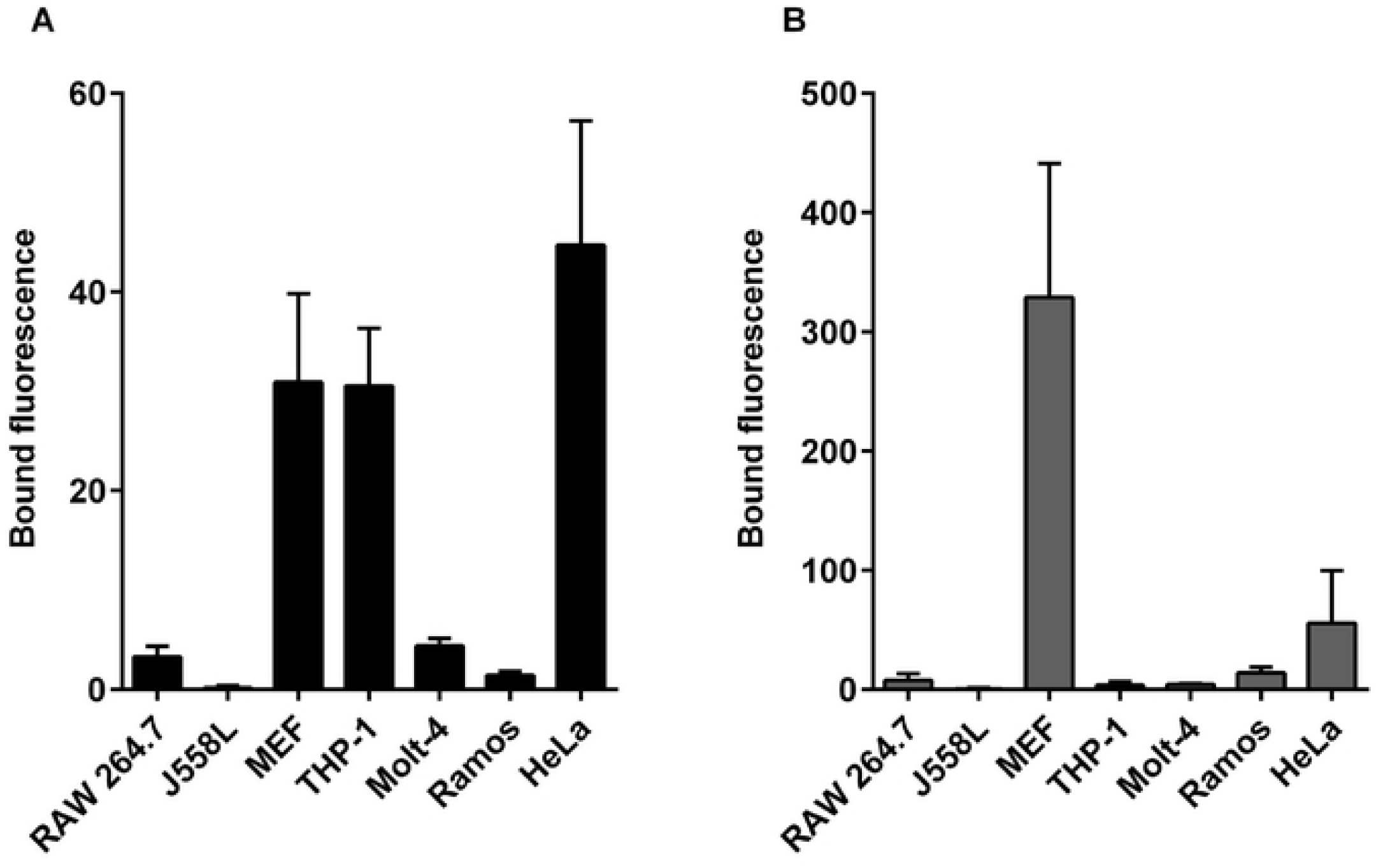
Binding of labelled GST-CNFY to immune cells. Indicated cell lines were analyzed for binding of DL488-GST-CNFY or a FITC-labeled antibody against HSPG by flow cytometry. Given is the average of bound fluorescence ± SD (n *=* 3) for binding **(A)** of DL488-GST-CNFY (black) and **(B)** a FITC-HSPG antibody (grey).

### Characterization of a possible CNFY-GAG interaction

We previously showed, that sodium chlorate, which inhibits sulfation of GAG chains, without affecting the size of the sugar chains, reduces intoxication of HeLa cells by CNFY, indicating that proteoglycans may be relevant for toxin binding (10, 15). Following data from the human protein atlas, proteoglycans are widely expressed by most mammalian cells, however, circulating blood cells only marginally express them (Fig. S1A). Therefore, we studied expression of GAG on the same cell lines analyzed above. Cells were incubated with a FITC coupled antibody detecting HSPG and washed. Bound fluorescence was then analyzed by FACS measurements. In line with toxin binding, MEF and HeLa cells bound the antibody (Fig. 1B). Surprisingly, there was no bound fluorescence detectable on THP-1 cells, suggesting that this cell line may not express HSPG and also other PG (for example chondroitin sulfate or dermatan sulfate) may bind the toxin. In Molt-4 cells (human T-lymphocytes) most of membrane GAG like syndecans and glypicans were not expressed (Fig. S1A). Only glypican-2 RNA was detected more prominently in Molt-4 and THP-1 compared to HeLa cells. If poly-sugars like GAG are involved in toxin interaction, binding kinetics should show no saturation. Therefore, we incubated HeLa cells and THP-1 cells, respectively with increasing concentrations of DL488-GST-CNFY. As shown in Fig. 2A, increasing concentrations of the protein led to increased binding without obvious saturation, indicating a highly abundant structure (like sugars) present on mammalian cells as interaction partner of the toxin. However, data fit best to regression with two curves, one saturated, most likely a protein receptor, the other one not saturated, most likely a sugar. This suggests a combined function of at least two binding surfaces on the cell (Fig. S2). To study which part of the toxin may interact with GAG, we repeated the experiments with the labeled C-terminal domain (GST-CNFY_709-1014_), because it was shown for the homologue toxin family member CNF1 that this domain mediates binding to its receptor Lu/BCAM (16). As for the full toxin, we could not observe saturated binding of GST-CNFY_709-1014_ to HeLa cells, indicating that the C-terminal part of CNFY is sufficient for GAG binding (Fig. 2A). As expected, competition experiments on HeLa cells with DL488-GST-CNFY and unlabeled GST-CNFY decreased binding of the labeled full toxin. However, competition with the C-terminal domain (GST-CNFY_709-1014_) did not influence binding of the labeled full toxin. In line with the mathematical curve fits (Fig. S2) this suggests that different cellular structures add to toxin binding (Fig. 2B). To directly study the involvement of GAG on the cell surface in CNFY binding, we specifically removed HS-chains from GAG by treating cells with a recombinant active human heparanase 1 (HPSE). Therefore, HeLa cells were incubated with active HPSE, washed and analyzed for toxin binding. As control for HPSE activity, we studied also binding of an anti HSPG antibody to the cells. As shown in Fig. 2C, treatment of HeLa cells with HPSE led to reduced binding of labeled GST-CNFY. Binding of GST-CNFY_709-1014_ was even slightly stronger inhibited (by about 40%) equal to the reduced binding of a FITC-labeled anti HSPG antibody (Fig. 2C). The data indicate that HSPGs are involved in CNFY (more specifically CNFY_709-1014_) binding to cells.

**Figure 2:**
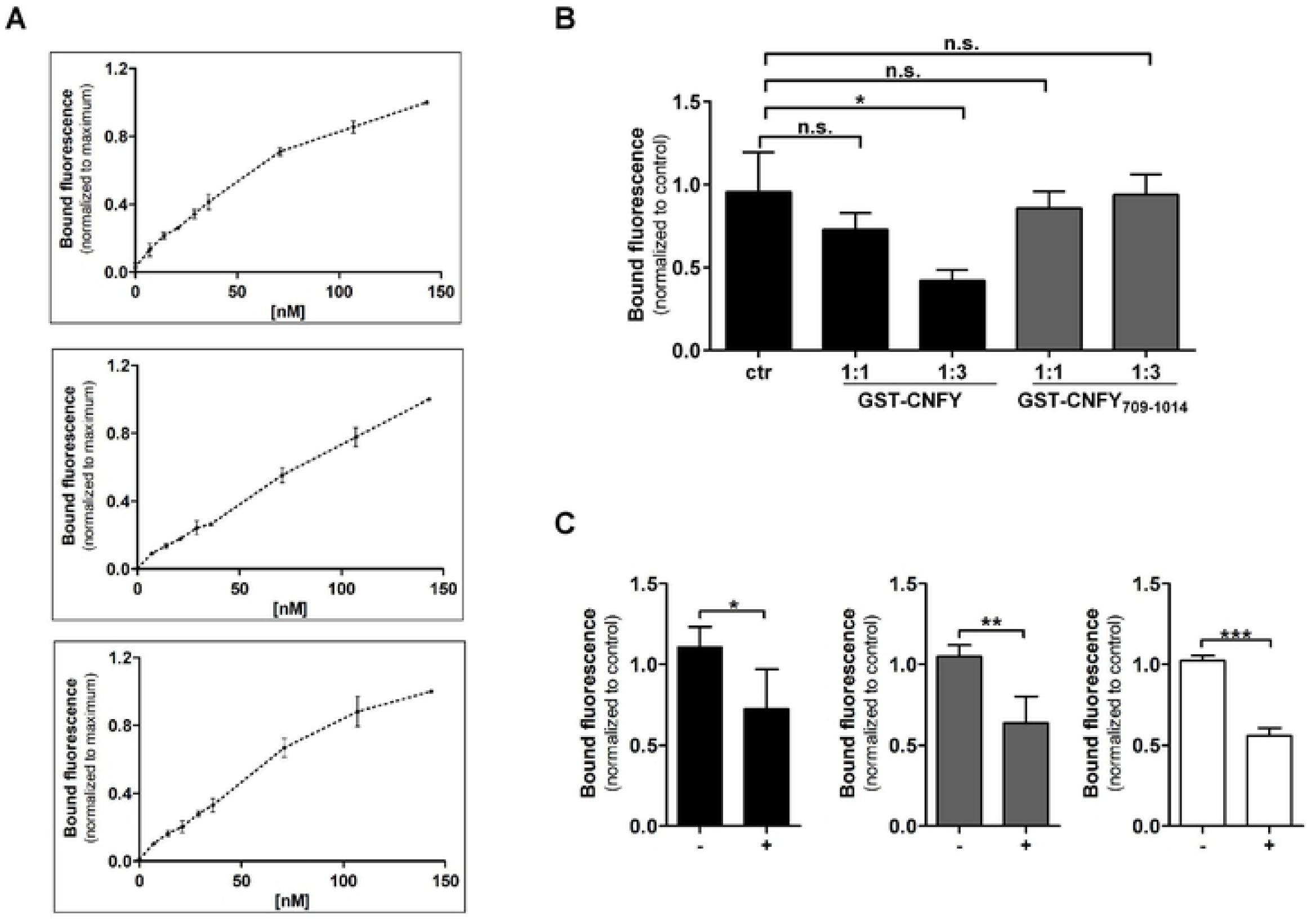
Proteoglycans are involved in CNFY binding. **(A)** HeLa or THP-1 cells (middle) were incubated with increasing concentrations of DL488-GST-CNFY (left, middle) and DL488-GST-CNFY_709-1014_ (right), respectively. Bound fluorescence was measured by flow cytometry. Data were normalized to the detection at the highest toxin concentration and represent the mean ± SD of three independent experiments. **(B)** HeLa cells were co-incubated with DL488-GST-CNFY, and unlabeled GST-CNF or GST-CNFY_709-1014_ as indicated and washed. Bound fluorescence was analyzed by flow cytometry. Given is the average of bound fluorescence ± SD normalized to control cells only incubated with DL488-GST-CNFY (n *=* 2). One-way ANOVA was applied for statistical comparison. ** p < 0.01. **(C)** Flow cytometry analyses of HeLa cells pre-incubated or not with activated human Heparanase-1 (HPSE) before DL488-GST-CNF proteins (GST-CNFY, (black bars) and GST-CNFY_709-1014_ (grey bars) or a FITC-labeled antibody against HSPG (white bars) were added, respectively. Results are representatives of 3 independent experiments shown as mean ± SD. Statistical analyses were performed using student’s t-test. Non-significant (n.s.); * p < 0.05 ** p < 0.01 *** p < 0.001.

### Identification of the relevant GAG

To study which GAG might be involved in CNFY binding, we carried out competition studies with several soluble GAG. To this end, HeLa cells were incubated with DL488-GST-CNF proteins in the presence and absence of, heparin (H), chondroitin sulfate A (CS), dermatan sulfate (DS), hyaluronic acid (HA), dextran sulfate (DxS) and fondaparinux (F) (Fig. 3A). As a control, also binding of labeled *Clostridium difficile* toxin A (TcdA) was studied. It was recently shown that TcdA binds to sulfated GAG (14). In the presence of heparin, hyaluronic acid and dextran sulfate, respectively, there was reduced binding of TcdA to Hela cells (Fig. 3B). Also binding of CNFY full length toxin was diminished by addition of dextran sulfate. Importantly, there was nearly complete inhibition of binding of CNFY_709-1014_ in the presence of heparin, indicating that CNFY C-terminal domain has high affinity to heparin and presumably binds to heparan sulfate HSPG on the cell surface, which is in line with the results shown above (using heparanase 1). Heparin, dermatan sulfate and dextran sulfate showed an inhibitory effect on CNFY_709-1014_. However, reduced binding of the full toxin was only marginally inhibited (significant only with dextran sulfate) and was not sufficient, to block intoxication of cells. RhoA was modified in all cells treated with CNFY or CNF1, respectively, independent of the competitors added. As expected, the catalytically inactive mutants of both toxins (C866S) did not induce a shift of RhoA (Fig. S3). Importantly, the reduced amount of toxin bound to the cell surface seems still sufficient to modify the complete pool of RhoA within the cells. Taken together, the isolated C-terminal domain of CNFY interacts strongly with heparin and binds to dermatan and dextran sulfate. Binding of the full toxin was not influenced dramatically in the presence of polysugars.

**Figure 3:**
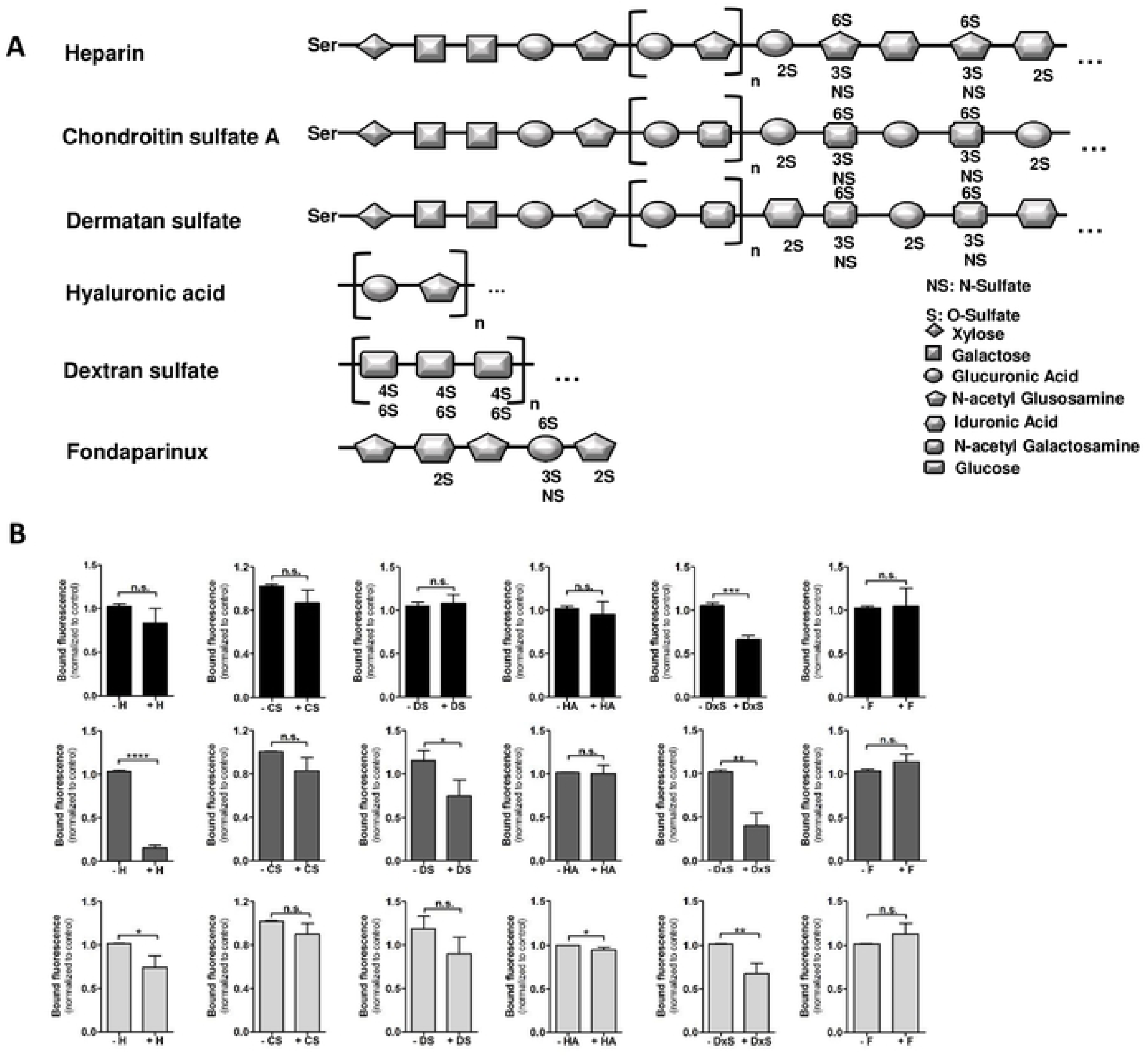
Characterization of the CNFY-GAG interaction. **(A)** Simplified scheme of glycosaminoglycans: Symbols used are Ser: Serine, 2S: 2-*O*-sulfated, 3S: 3-*O*-sulfated, 6S: 6-*O*-sulfated. **(B)** For flow cytometry HeLa cells were incubated with different DL488-GST-CNF proteins (GST-CNFY (black), GST-CNFY_709-1014_ (grey) or His-TcdA (light grey)), in the presence or absence of GAG: heparin (H), chondroitin sulfate A (CS), dermatan sulfate (DS), hyaluronic acid (HA), dextran sulfate (DxS) and the pharmacon fondaparinux (F). Data shown represent the mean of the bound fluorescence ± SD (n = 3). Statistical analysis was performed using student’s t-test. Non-significant (n.s.); * p < 0.05; ** p < 0.01; *** p < 0.001; **** p < 0.0001.

### Direct interaction of CNFY_709-1014_ with heparin

To further characterize the binding affinities of CNFY_709-1014_ with heparin, we utilized the bio-layer interferometry (BLI) assay. Binding of a protein to immobilized heparin results in a shift within the light interference pattern that can be monitored in real-time. The catalytic inactive full toxin CNFY_C866S_ was used as binding control, whereas biotinylated hyaluronic acid and biotinylated cellulose served as sugar control. As shown in Fig. 4A, the full toxin only marginally interacted with heparin and did not bind to hyaluronic acid or cellulose, as expected from the competition studies (Fig. 3B). In contrast, there was strong binding of CNFY_709-1014_ to heparin but no interaction to the sugar controls (Fig. 4B). The homologue C-terminal domain of the *E*.*coli* toxin CNF1 (CNF1_709-1014_, negative control) only weakly interacted with all studied sugars (Fig. 4C). Affinity of CNFY_C866S_ and CNFY_709-1014_ was analyzed measuring binding curves with increasing concentrations of heparin (0.1 to 1 mg/ml, Fig. 4D, E). The C-terminal domain of CNFY has high affinity to heparin with a K_D_ in the low µmolar range (about 10 µM). In contrast, the affinity of the full CNFY to heparin was too low to estimate a K_D_. CNFY_709-1014_ contains a typical heparin binding motif rich in lysine (^772^VKKTKF^779^). To analyze its potential involvement in GAG binding, we mutated each lysine in this motif to alanine (structure in S4A). We then labeled the respective recombinant proteins (CNFY_709-1014_) with DL488 and studied binding to HeLa cells in the presence and absence of heparin. All mutants bound to HeLa cells. Moreover, in all cases heparin diminished the amount of surface-bound toxin. However, inhibition of CNFY_709-1014_ K774A binding by heparin was significantly reduced (Fig. S4B). Moreover, BLI assays revealed that binding of CNFY_709-1014_ K774A to immobilized heparin was weaker compared to wildtype CNFY_709-1014_ (Fig. S4C).

**Figure 4:**
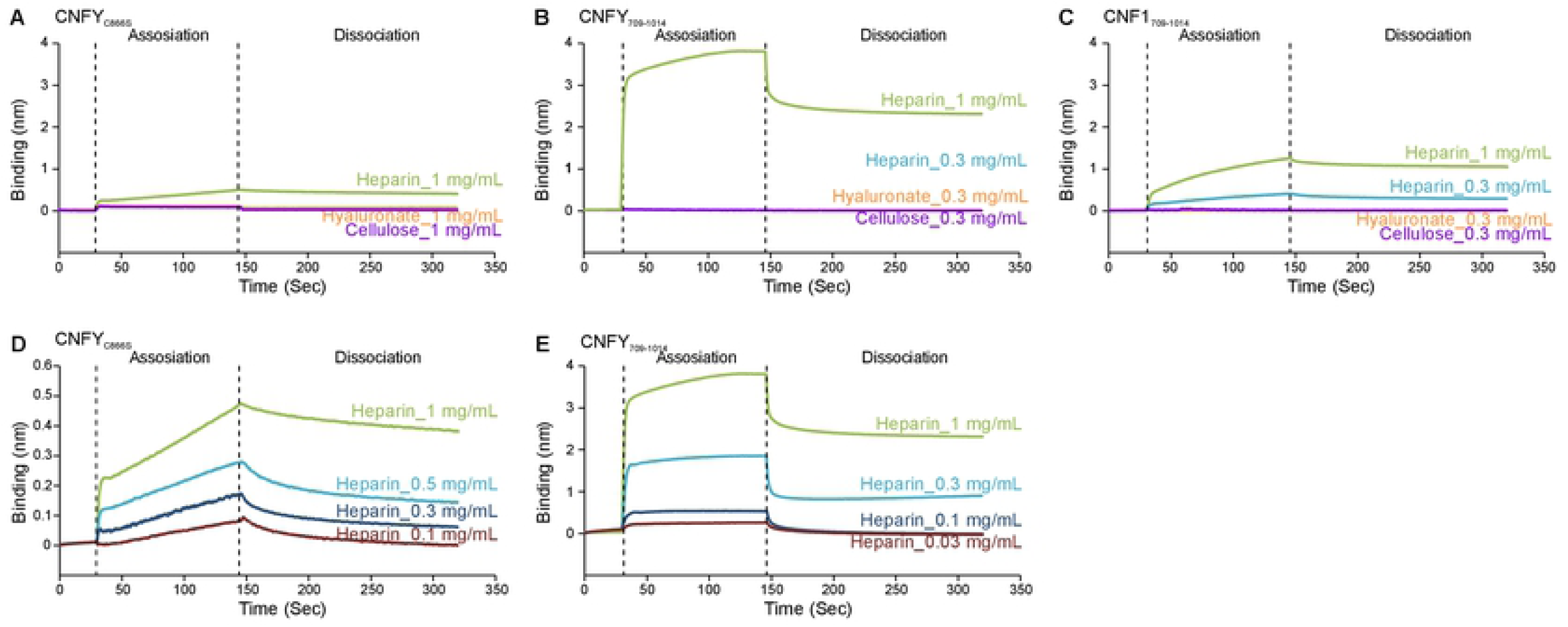
Direct interaction of CNFY_709-1014_ with heparin. The binding affinity to immobilized biotin-heparin was analyzed using BLI assays. Biotin-hyaluronic acid and biotin-cellulose served as sugar controls. Representative data was shown from two independent experiments for **(A, D)** CNFY_C866S_, **(B, E)** CNFY709-1014, **(C)** CNF1709-1014.

### GAG are cellular attachment factors for CNFY but not sufficient for intoxication

To study the possible involvement of GAG in toxin binding and uptake, we made use of a recently generated knockout cell line of Chinese hamster ovary (CHO) cells. These cells do not express Xylose transferase 1, the enzyme transferring the first sugar (Xylose) to the acceptor serine of the PG core proteins (Fig. 5A). Although CHO cells express high amounts of GAG, FACS experiments with labeled toxins showed that CNFY did not bind to XYLT^-/-^ cells nor to CHO wildtype cells. The isolated C-terminal domain of the toxin (CNFY_709-1014_) still interacted with CHO cells whereas binding to XYLT^-/-^ cells was nearly completely blocked (5B). As shown in Fig. 5C, CHO wildtype and XYLT^-/-^ cells are not intoxicated with CNFY (GST-CNF1 was used as positive control), which is in line with the binding studies. This indicates that the isolated C-terminal domain is more accessible for GAG binding and the binding site may be sterically hidden in the full toxin. Since a conformational change occurs following acidification within the endosome, sugar binding of the CNFY C-terminal domain may play a role within the endosome rather than on the cell surface. To be able to distinguish the three steps required for intoxication: binding to cells, endocytosis and release into the cytosol, we made use of a second cellular model based on HeLa cells and deficient in the glucosyltransferase Exostosin-2 (EXT2, (17)). These cells express GAG, which carry shortened heparan sulfate chains, because EXT2 is specifically required for the elongation of the HS chain and no other types of GAG (12). FACS experiments with these cells showed 72% reduced binding of full length CNFY, as well as for CNFY_709-1014_, about 87% reduction (Fig. 5D), respectively, compared to wildtype HeLa cells. As a control for the functional knockout, we used TcdA, as well as a FITC-HSPG antibody. Also binding of TcdA and the HSPG-antibody to the EXT2^-/-^ cells was 70% respectively 87% reduced (Fig. 5D). The data show that heparan sulfate chains are involved in CNFY attachment to the cell surface, but they may not be the unique binding structures. We again studied intoxication of HeLa and EXT-2^-/-^ cells by analyzing the shift of deamidated RhoA in the lysates of toxin-treated cells. However, although we measured binding of CNFY to EXT2^-/-^ cells, no modification of RhoA was detectable (Fig. 5E). Interestingly, these data suggest that the toxin did bind to EXT2^-/-^ cells but was not able to enter the cytosol.

**Figure 5:**
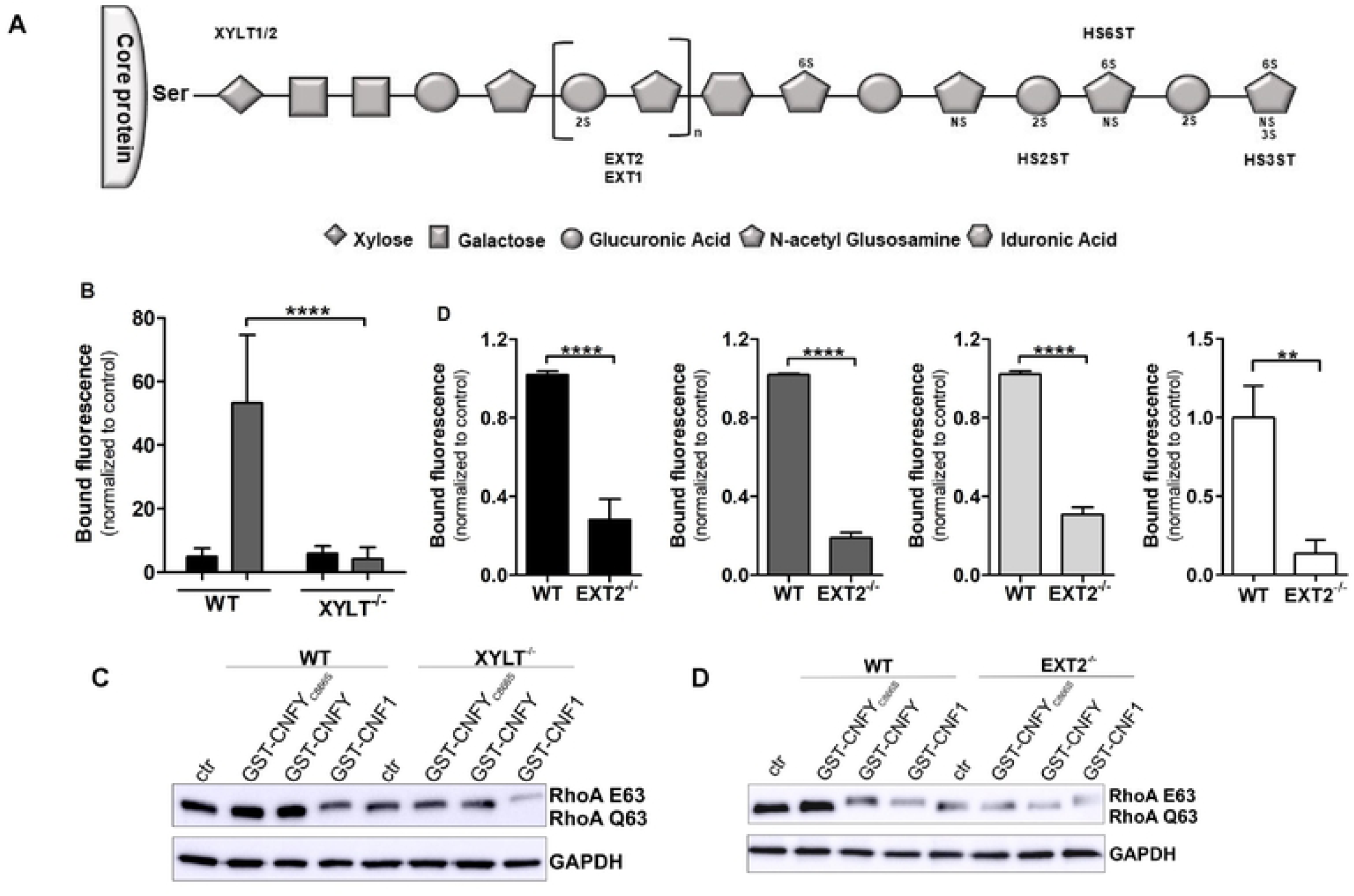
GAGs are cellular attachment factors for CNFY but not a crucial membrane receptor. **(A)** Simplified scheme of different enzymes involved in HS biosynthesis (enzymes gene names: XYLT1/2: Xylosyltransferase 1/2, EXT1/2: Exostosin Glycosyltransferase 1/2, HS6ST: Heparan Sulfate 6-O-Sulfotransferase 1, HS3ST: Heparan Sulfate-Glucosamine 3-Sulfotransferase 1). The symbols used for the saccharide units are defined below the scheme. Ser: Serin, NS: N-sulfated, 2S: 2-O-sulfated, 3S: 3-O-sulfated, 6S: 6-O-sulfated. **(B)** Flow cytometry analysis of DL488-GST-CNFs (GST-CNFY (black), GST-CNFY_709-_ 1014 (grey)) binding to CHO wildtype (WT) and CHO Xylosyltransferase knockout (XYLT^-/-^). Given is the average of bound fluorescence ± SD (n *=* 2). Statistical analysis was performed using one-way ANOVA. **** p < 0.0001. **(C)** Shift assay showing deamidated RhoA (E63). CHO WT and CHO XYLT^-/-^ cells were intoxicated with GST-CNFY, GST-CNF1 or the catalytic inactive mutant (GST-CNFY_C866S_) and lysed. Deamidation of RhoA was detected by Western blotting using an anti-human RhoA antibody. Equal loading was verified by detection of GAPDH. The experiments were repeated three times independently with similar results. **(D)** Bound fluorescence was measured in HeLa wildtype (WT) and Exostosin-2 knockout (EXT2^-/-^) cells by flow cytometry after incubation with labeled toxins: GST-CNFY (black), GST-CNFY_709-1014_, TcdA (light grey) or with a FITC-HSPG antibody (white). Data shown represent three independent experiments, the mean of the bound fluorescence ± SD. Statistical analyses were performed using one-way ANOVA. *** p < 0.001; **** p < 0.0001. **(E)** Shift assay showing deamidated RhoA (E63). HeLa wildtype (WT) and Exostosin-2 knockout (EXT2^-/-^) cells were incubated with GST-CNFY, GST-CNF1 or the catalytic inactive mutant (GST-CNFY_C866S_) and lysed. Cell lysates were analyzed by Western blotting using an anti-human RhoA antibody. GAPDH was used as loading control. The experiments were repeated three times independently with similar results.

### CNFY as well as CNFY_709-1014_ are enriched in early endosomes

Importantly, the C-terminal part of the toxin (CNFY_709-1014_) comprises the catalytic domain. Therefore, it is surprising that this domain binds with high affinity to heparin and heparan sulfate side chains whereas the full toxin did not. We asked what function sugar binding to the catalytic domain of a toxin could hold and where this interaction might occur. We studied, whether the isolated catalytic domain is sufficient to be taken up by endocytosis. Therefore, we analyzed localization of labeled GST-CNFY full and GST-CNFY_709-1014_, respectively following incubation of HeLa cells by co-staining with an antibody against Rab5, a marker for early endosomes. As shown in Fig. 6, both, the full toxin and the C-terminal domain co-localize with Rab5 and therefore appear enriched in early endosomes. This indicates that the interaction of CNFY_709-1014_ with HSPG is sufficient for endocytosis.

**Figure 6:**
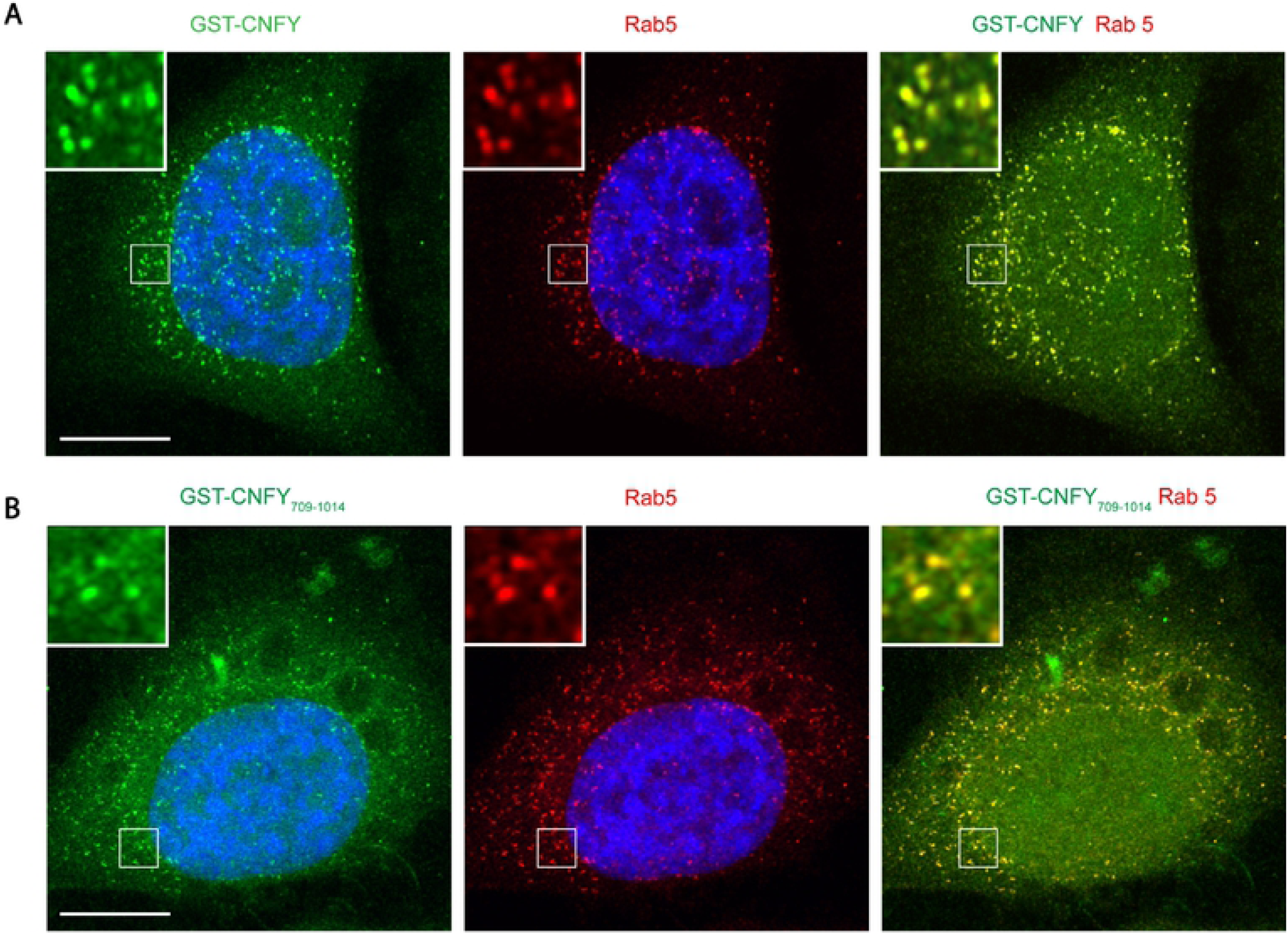
CNFY as well as CNFY_709-1014_ are enriched in early endosomes. Immunofluorescence analysis of the endosomal uptake of GST-CNFY **(A)** and GST-CNFY_709-1014_ **(B)** labeled with Dyelight488 (green) after 30 min incubation with HeLa cells. Early endosoms were stained in red with a Rab5 antibody. Cell nuclei were labeled with DAPI. Scale bar represents 10 μm.

Together our data suggest that CNFY binding to cells with shortened sugar chains on the surface GAG is reduced but not completely missing. However, intoxication of the cells, which requires endocytosis and release from the acidified endosomes is eventually blocked, suggesting a specific function of GAG within the endosome.

### Heparin interaction with CNFY does not induce a conformational shift

To answer the question whether the shift to acidic pH or the interaction of HSPG with CNFY induces a conformational change, we measured the distance between the two single cysteine residues (C134 and C866) in CNFY using nitroxide spin labeling in combination with PELDOR spectroscopy (18). The PELDOR method exploits the distance dependence of the interaction between the magnetic dipole moments of two spin labels. Nitroxide spin-labeled CNFY samples were frozen and subjected to a four-pulse PELDOR sequence. The resulting echo decays were first baseline corrected (Fig. 7, blue curves), then analyzed for dipolar oscillations, and their frequencies correlated to distance distributions (Fig. 7, black curves) between the points of highest electron spin-density in the two nitroxide radicals covalently attached to C134 and C866. Prominent peaks in the distance distributions can be observed between 4 and 5 nm. Specifically, CNFY at physiological pH 7.4 exhibits a peak distance of 4.1 nm (Fig. 7A), which shifts to 4.7 nm upon lowering the pH to 5 (Fig. 7C). Addition of heparin did not alter these values within the experimental error (Fig. 7, panels B and D). Hence, PELDOR data indicate that the acidic pH rather than heparin binding induces a conformational change of the toxin.

**Figure 7:**
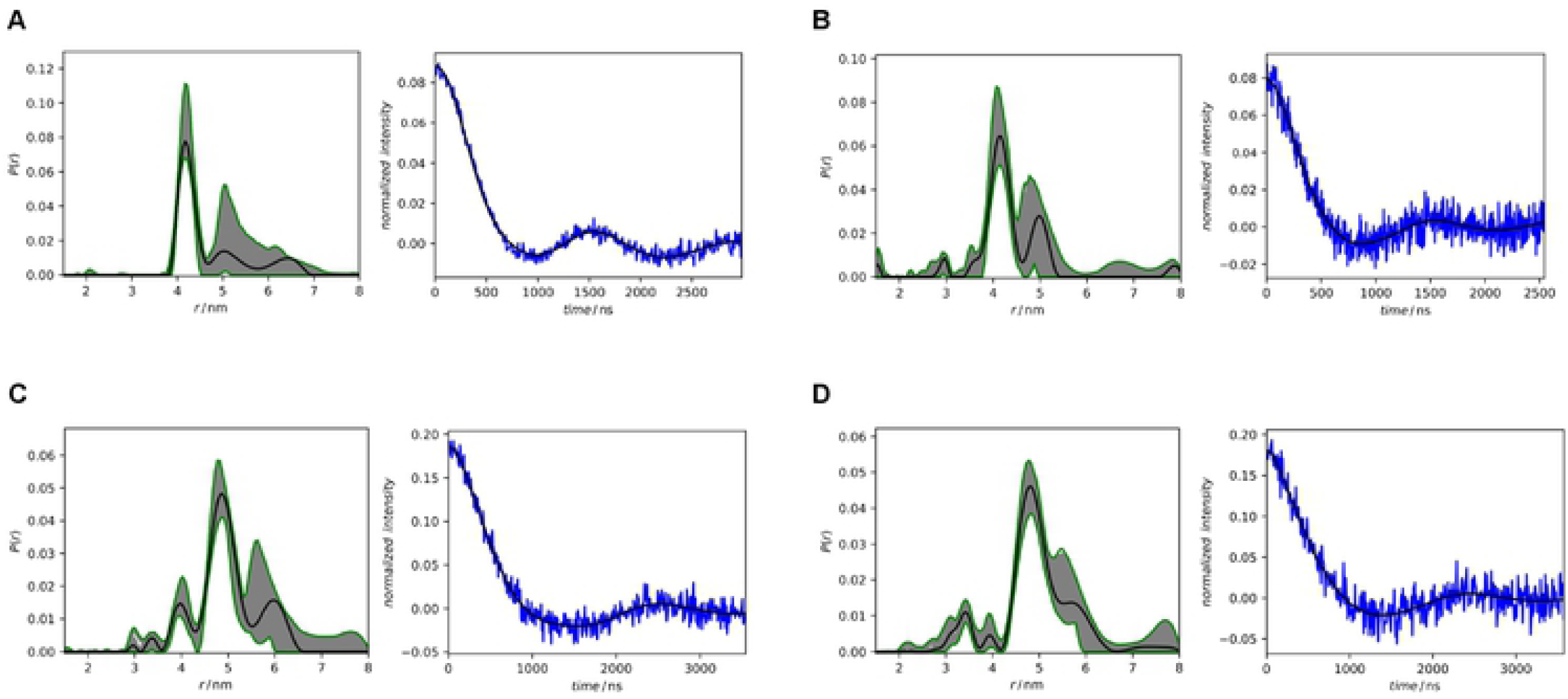
Heparin interaction with CNFY does not induce a conformational shift. Baseline-corrected PELDOR time-domain data (blue curves) together with fit functions (superimposed black solid lines) and calculated distance distributions (black solid lines) with validation range (grey-shaded areas) of nitroxide-spin labeled CNFY at physiological pH (A, B) and acidified (C, D). Panels B and D show CNFY samples that were supplemented with 50µg/ml heparin.

### Interaction of CNFY with heparin is unaffected by the pH

Because the acidic pH of endosomes is required for toxin release, which was previously shown by bafilomycin A1, an inhibitor of the endosomal proton pump (1, 16), we studied whether heparin interaction with CNFY is pH dependent. Therefore, we acidified the cell culture medium to mimic the pH present in late endosomes on the surface of HeLa cells. Under these conditions competition assays with heparin were performed.

Therefore, HeLa cells were incubated with the labeled CNFY or labeled CNFY_709-1014_ at pH 7.4 or pH 5.0 in the presence or absence of heparin, respectively as indicated in Fig. 8. In the absence of heparin, binding of labeled CNFY to the cells was slightly diminished at acidic pH. A further inhibition by heparin was visible, but not significant (Fig. 8A). In contrast, binding of labeled CNFY_709-1014_ was even increased at acidic pH but significantly inhibited in the presence of heparin at acidic and neutral pH, respectively, indicating that binding of the C-terminal part of CNFY to heparin is not diminished at acidic pH (Fig. 8B). Acidification of the endosomes seems not sufficient to separate the toxins C-terminal domain from membrane bound HSPG. Membrane interaction is required for the passage through the membrane. However, coming off the membrane should be a prerequisite for the toxins release into the cytosol.

**Figure 8:**
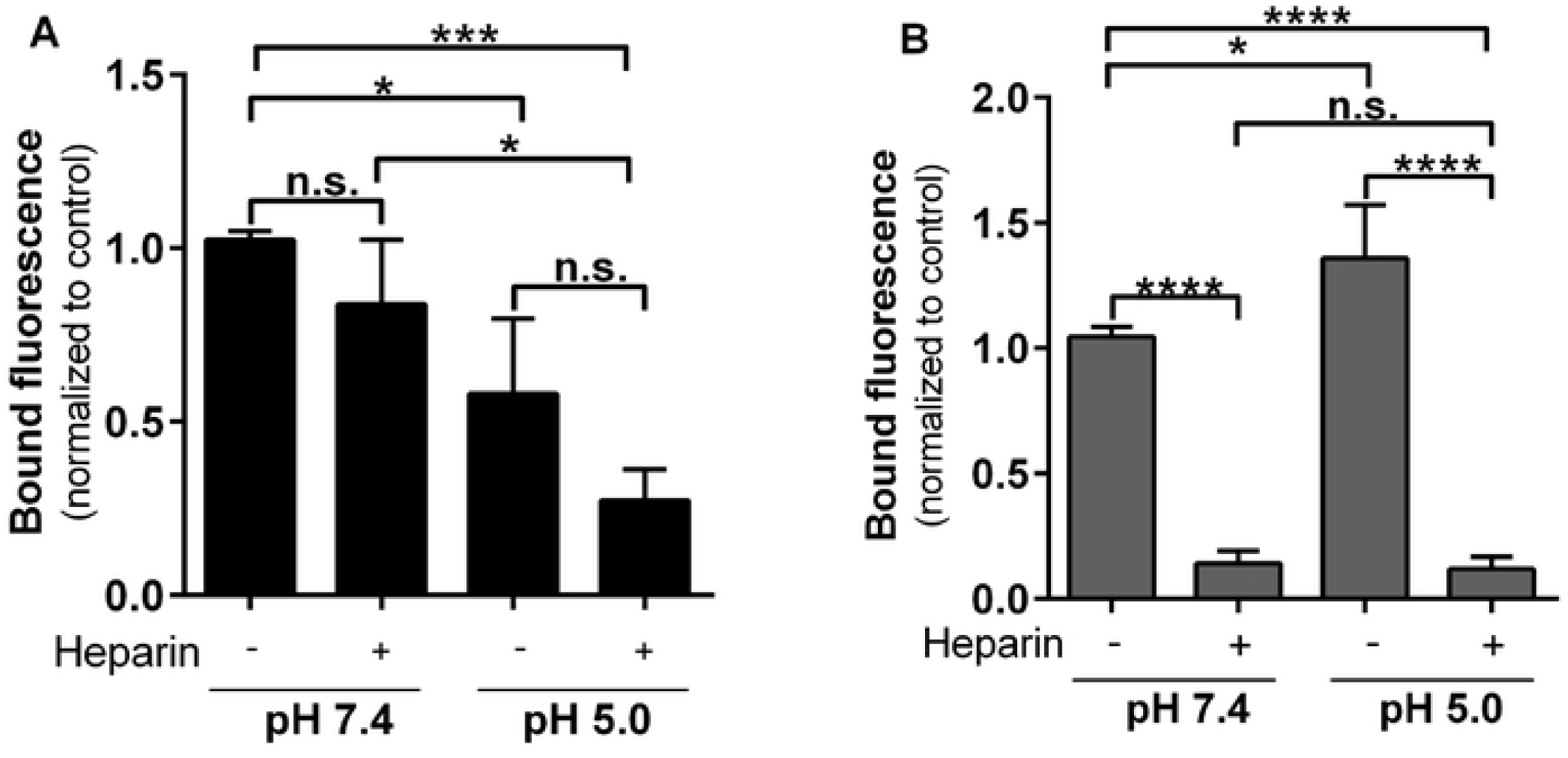
Interaction of CNFY with heparin occurs at neutral and acidic pH. HeLa cells were co-incubated with different DL488-labeled toxins: GST-CNFY (black) or GST-CNFY_709-1014_ (grey) in absence or presence of Heparin at pH 7.4 or pH 5.0, respectively as indicated. Binding of DL488-GST-CNF proteins without Heparin at pH 7.4 serves as a control. Bound fluorescence was measured by flow cytometry. Data shown represent three independent experiments, their mean of the fluorescence ± SD. Statistical analyses were performed using one-way ANOVA. Non-significant (n.s.); * p < 0.05; *** p < 0.001; **** p < 0.0001.

### Cleavage of heparin is required for the release of CNFY from the endosome

High affinity binding of the C-terminal domain to heparan sulfate within the endosome would interfere with its release into the cytosol. It is well known that an acidic pH not only induces a conformational change of proteins and activates proteases but also leads to higher activity of heparanases (19). Cleavage of heparin would also relieve the toxin from its attachment site at the inner endosomal membrane. Therefore, we intended to inhibit the activity of heparanase by reducing its expression with siRNA and to study its effect on cell intoxication. As shown by Western Blotting, siRNA significantly reduced the amount of heparanase in HeLa cells (Fig. 9A, B). In line with the assumption of a crucial role of heparinase in the intoxication process, deamidation of RhoA in siRNA-treated cells occurs slower compared to cells treated with control siRNA (Fig. 9C).

**Figure 9:**
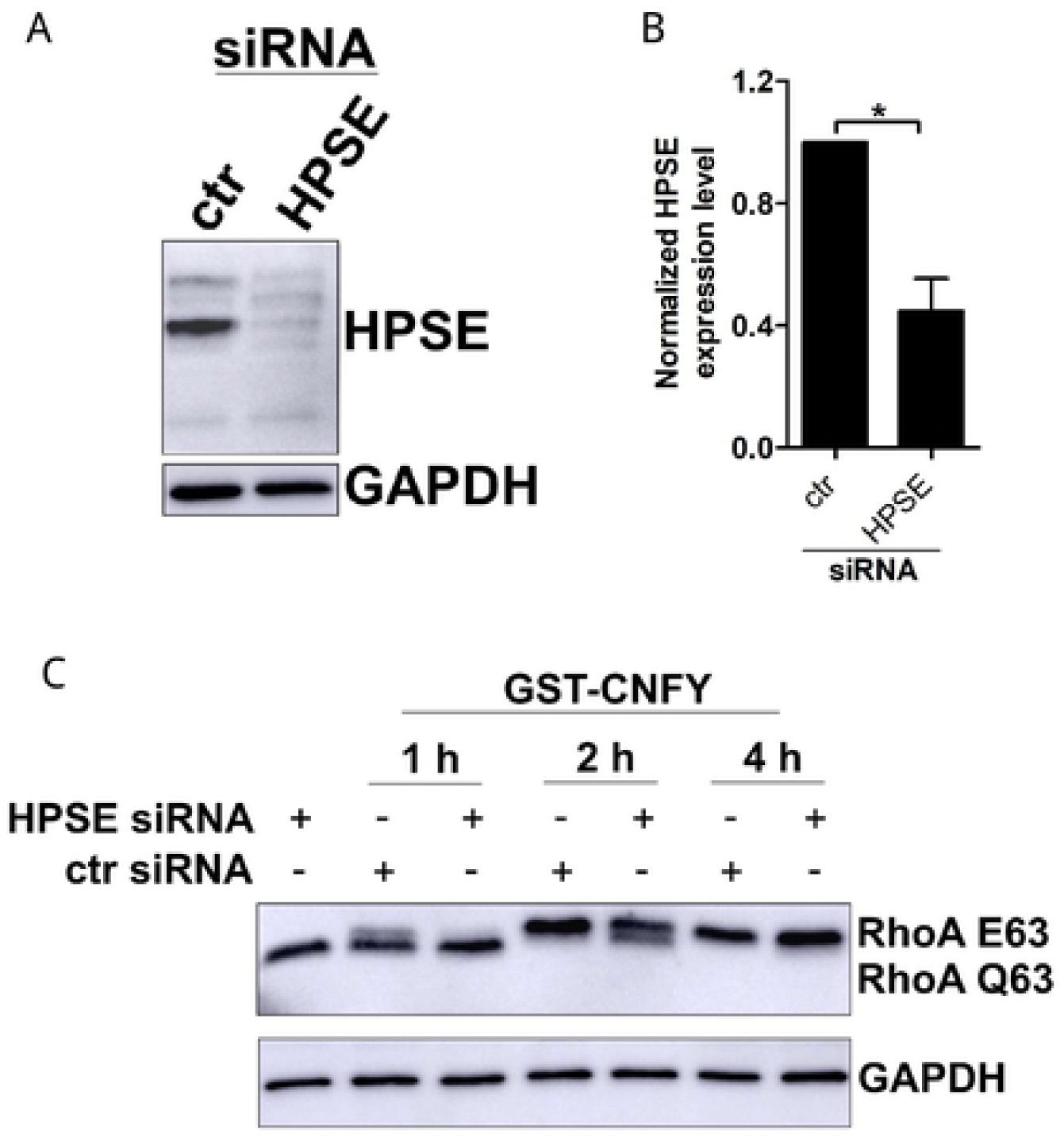
Cleavage of heparin is crucial for the release of CNFY from the endosome. **(A)** SiRNA based knockdown of HPSE was confirmed by Western blotting using an anti-human HPSE 1 antibody. **(B)** Quantification of three independent experiments shown in A given as mean ± SD. Statistical analysis was performed using student’s t-test. * p < 0.05. **(C)** Time dependent shift assay: SiRNA-treated HeLa cells were incubated with GST-CNFY for 1-4 h. RhoA was detected by Western blotting following urea SDS PAGE. GAPDH was used as a loading control. The experiment was repeated three times independently with similar results.

### Model for binding and uptake of CNFY

Our data lead to the following model for the cellular uptake of CNFY (Fig. 10). CNFY binds with its N-terminal domain to an unknown protein receptor, which may already lead to a conformational change of the toxin allowing an interaction of the C-terminal domain with HSPGs thus enhancing avidity. Following receptor-mediated endocytosis and acidification, a conformational change of CNFY leads to cleavage retaining high affinity contact of the C-terminal domain to HS side chains at the membrane. Acidification allows incorporation of the translocation domain into the endosomal membrane and leads to activation of heparanase. The C-terminal part of CNFY would remain attached to the inner endosomal membrane by its high affinity interaction to HS. Release into the cytosol therefore requires cleavage of HS by the activated heparanase.

**Figure 10:**
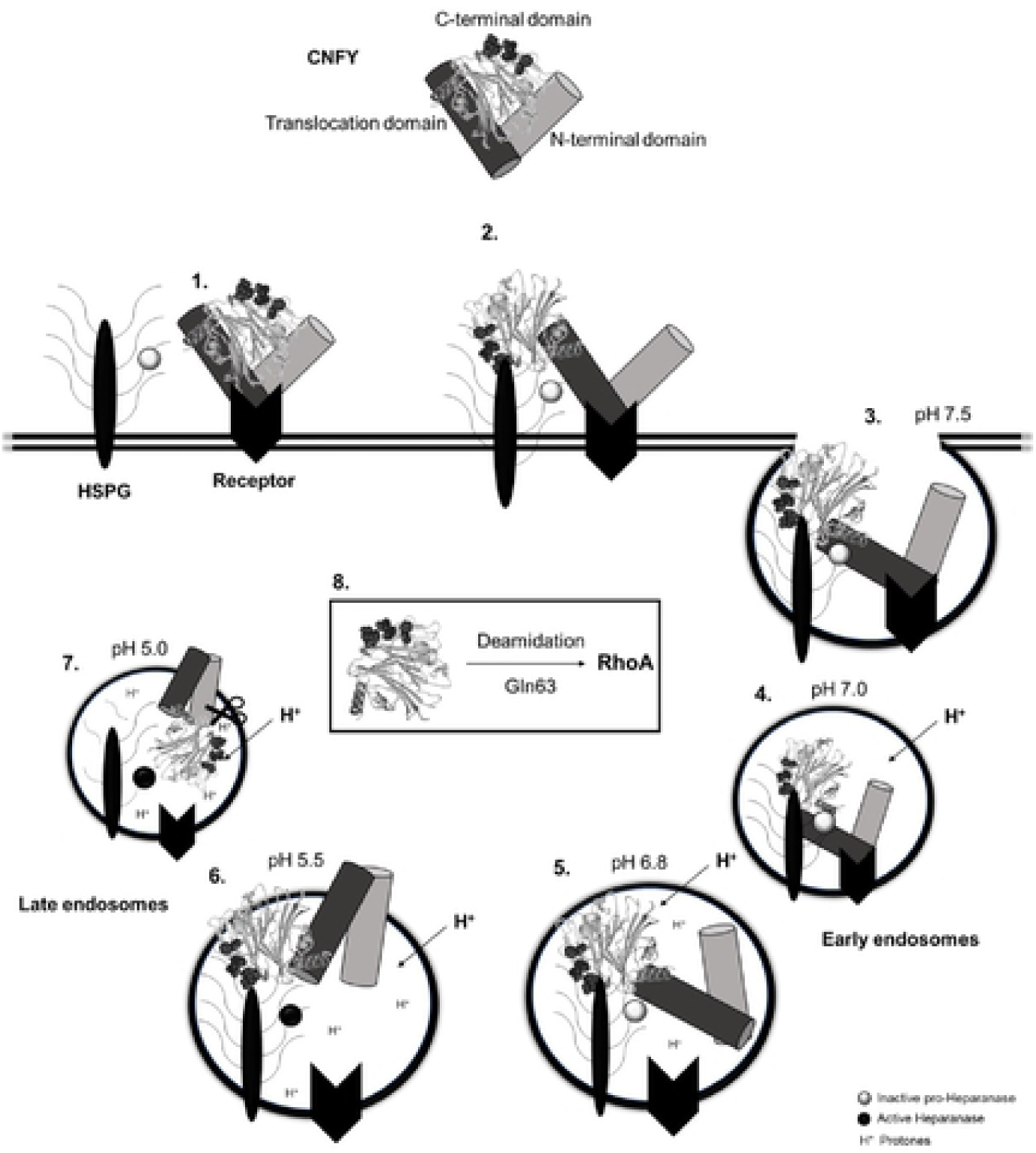
Cellular uptake of CNFY. (1) CNFY binds to a protein receptor on the host cell membrane. (2) Binding of CNFY to the protein receptor induces a conformational change which allows the interaction of the c-terminal domain to heparansulfates (HS). (3) The CNFY-receptor complex as well as the bound inactive pro-heparanase is taken up via receptor-mediated endocytosis. (4+5) Early endosomes are acidified due to a vacuolar H+-ATPase leading to dissociation of the toxin-receptor complex, whereas the interaction to HS is still existent. (6+7) The acidic pH of the late endosome induces partial unfolding of the toxin provoking the exposure of the hydrophobic regions within the translocation domain eventually allowing incorporation into the endosomal membrane. In contrast to the toxin-protein receptor interaction, binding of the c-terminal part to HS even increases at pH 5.0, due to a more positive net charge of the protein. (8) Instead, the c-terminal domain is released into the cytosol by the activated heparanase due to cleavage of the sugar chains. In the cytosol, the toxin deamidates RhoA at glutamine 63, leading to constitutive activation of the GTPase.

## Discussion

In a recent publication by Heine et al., a relevant influence of CNFY on pro-inflammatory IL-6 production in mice was reported. Surprisingly, binding and intoxication studies indicated that CNFY does not influence immune cells directly. IL-6 is expressed by immune cells but also by fibroblasts and endothelial cells. Following our results presented here, the later are more likely targets of CNFY. Based on these findings, we intended to study the reason for the missing immune cell response by analyzing binding and uptake of CNFY in more detail. Our group showed before, that treatment of cells with sodium chlorate, which induces under-sulfation of cellular proteoglycans, inhibits intoxication of HeLa cells (10), indicating that sulfated sugars, which are rarely present on immune cells, could be involved in a crucial step of the toxins entry path. Interestingly, it has been shown that heparan sulfate proteoglycan (HSPG) binding of viruses allows movement on the cell surface until the protein receptor is bound to mediate endocytosis (20). To verify our former results, we directly removed HSPG form the cell surface by heparanase treatment. A specific antibody recognizing an epitope of heparan sulfate (21) revealed a 50% reduction of HSPG. Binding of labeled GST-CNFY or GST-CNFY_709-1014_, respectively was reduced to a similar degree. Binding experiments with increasing amounts of toxin suggested combined interaction of CNFY with a protein and an abundant surface molecule like a sugar as binding partner. Our FACS data fit best to regression with two curves, one saturated, most likely a protein receptor with high affinity (Fig. S2, blue curve), the other one additive and not saturated (Fig. S2, green curve). Following this assumption, the linear, not saturated curve would start at the brake point concentration of about 25 to 30 nM. However, with two binding partners of one protein also avidity besides affinity would define the binding kinetics. Moreover, interaction with surface sugars may not be determined by a single binding site, because sugars appear as trees. In line with a sugar tree as highly abundant surface interaction partner, binding curves of CNFY to cells showed no saturation. However, other structures present in high amounts are also present on the cell surface, for example lipids. Pore forming toxins like *Clostridium perfringens* alpha toxin bind to sphingomyelin (22), *Streptococcus pneumonia* pneumolysin binds with high affinity to cholesterol (23). To show toxin - sugar interaction more directly, competition studies with GAG were performed. Our data revealed that heparin and slightly weaker also dextran sulfate interfered with binding of labeled CNFY_709-1014_ to the cell surface. Binding of the full toxin was only marginally affected. Apart from dermatan sulfate, other polysugars like chondroitin sulfate, hyaluronic acid and the penta-sugar fondaparinux did not interfere with toxin binding.

The data show that the toxin-sugar interaction is not exclusively based on the charge but there seems to be a specific sugar-binding site. This is in contrast to the recently proposed binding of the clostridial toxin TcdA to the cell surface. The authors could interfere with intoxication of cells by addition of surfen (14). By comparison of several heparin interacting proteins, a conserved heparin binding motif built from basic amino acids (B) has been suggested, for example XBBXBX, XBBBXXBBBXXBBX or XBBBXXBX (25, 26).

With ^772^VKKTKF^779^ such a motif is present within the C-terminal domain. However, it is missing in CNF1, which is not influenced by Xylosyltransferase or Exostosin 2 knockout, respectively. Mutation of lysine774 to alanine in CNFY_709-1014_ reduced the inhibitory effect of heparin in cell binding experiments, indicating a crucial role of this amino acid for heparin binding. Exchange of K773 had no effect and mutation of K777 only marginally influenced heparin binding. Because the structure of CNFY can be modeled on the known structure of CNF1, we assume that the heparin-binding motif identified is located on the opposite side of the CNFY_709-1014_ substrate binding region. Therefore, heparin binding should not directly interfere with the catalytic activity of CNFY.

Although CHO cells do not bind and take up CNFY, they do bind the C-terminal part of the toxin, although interaction is weak. Therefore, CHO and the CHO-derived knockout cell line XYLT^-/-^ can be used to verify the specificity of the catalytic domain-sugar interaction by measuring binding of labeled CNFY_709-1014_. Cells deficient in Xylose transferase are not able to synthesize the surface GAG heparan sulfate, dermatan sulfate and chondroitin sulfate, two of which interfered with CNFY_709-1014_ binding to HeLa cells. As expected, compared to wildtype CHO-cells, binding of CNFY_709-1014_ to XYLT^-/-^ cells was significantly reduced to background levels, indicating a crucial role of heparan sulfate and dermatan sulfate for interaction with the C-terminal domain of CNFY. However, this interaction seems not to occur with the full toxin, suggesting a role for GAG binding in later steps of intoxication. A second explanation would be that the catalytic domain of the full toxin couldn’t interact with GAG because of sterical hindrance requiring a conformational change induced by protein receptor binding of the N-terminus. CHO cells were not intoxicated with CNFY, probably because of an absence of the protein receptor. Therefore, CHO-based cell lines can’t tell us about whether GAG binding is essential at any step of intoxication. To analyze the function of sugar binding in further steps of toxin uptake, we made use of a HeLa cell based knockout cell line: EXT2^-/-^ cells (14). Exostosin 2 is a glycosyltransferase, necessary for elongation of heparan sulfates. Without EXT2 the sugar chains appear drastically truncated. Binding of CNFY and CNFY_709-1014_ to EXT2^-/-^ cells was significantly reduced compared to wildtype HeLa cells, however not completely blocked. This indicates that CNFY interacts with GAG on cells expressing the receptor for the N-terminus. Together with the CHO binding studies these data support the hypothesis of a binding-induced conformational change allowing additional interaction with GAG and strengthening avidity. Most likely residual binding to EXT2^-/-^ cells was due to the presence of a second binding motif (protein receptor) or due to the presence of a similar sugar, like chondroitin sulfate. As shown in the GAG competition assays with HeLa cells, the residual toxin bound should be sufficient for intoxication of cells. However, no shift of deamidated RhoA was visible in lysates of toxin-treated EXT2^-/-^ cells, indicating that sugar binding may be crucial for a later step of intoxication: endocytosis or release from the endosome. Is HSPG a true receptor? Multiple functions of HSPGs are described: HSPGs are extracellular matrix components (27), they are found in secreted vesicles of mast cells (serglycin, (28)), described as sequestering co-receptors for growth factors and cytokines (fibroblast growth factor (FGF), (29, 30)) or binding partners for molecules undergoing transcellular transport (31). In principle, HSPG binding could be sufficient for endocytosis of the toxin. Therefore, we studied whether or not CNFY would be enriched within endosomes. Using microscopy, we found that endocytosis is not inhibited by knockout of Exostosin 2. However, no deamidation of RhoA occurred, indicating that the release into the cytosol is not possible. We suggest an additional function of HS binding within the endosome. Different from Diphtheria toxin, CNFs do not form a disulfide bond between receptor binding and catalytic domain. Cleavage would therefore release the catalytic domain into the lumen of the endosome and subsequent degradation. It has been shown, that acidification of endosomes is crucial for the release of many bacterial toxins into the cytosol. Bafilomycin A1, which inhibits acidification by blocking the endosomal proton pump, completely interferes with intoxication of HeLa cells (10). Acidification in the late endosomes leads to unfolding of the protein toxin, which allows insertion of two hydrophobic helices (located in the translocation domain) into the endosomal membrane. In PELDOR studies we showed that a conformational change of CNFY occurs following acidification, whereas there is no detectable change of conformation upon heparin binding. Moreover, heparin binding takes place at neutral and acidic pH. For CNF1 we showed that the protein chain is cleaved by a protease. This may also depend on the pH. The high affinity interaction to a membrane component should inhibit translocation from the endosome to the cytosol. It has not yet been shown that CNFY is cleaved by proteases. However, we assume that this is the case because of the high homology of the toxins (8). However, it is the C-terminal, catalytically active part, which shows affinity for heparin. Concerning our results presented here, a third function of acidification could be the activation of heparanase. This enzyme cleaves HS chains from HSPG within the endosome in a pH dependent manner. Recently, it was shown that acidification-dependent activation of heparanase is required for the release of a cytosol-penetrating antibody (19).

Knockdown of heparanase clearly diminished intoxication of HeLa cells with CNFY, indicating a crucial function of this enzyme for intoxication. Therefore, GAG binding of the catalytic domain of CNFY and cleavage of heparin seems to be crucial for intoxication. However, the deficiency of sugar chains should also lead to missing exposure of glycoproteins on the cell surface, among which the protein receptor may exist. We cannot study nor exclude this possibility, since the potential protein receptor is not yet known.

## Materials

Used antibody, application and supplier:

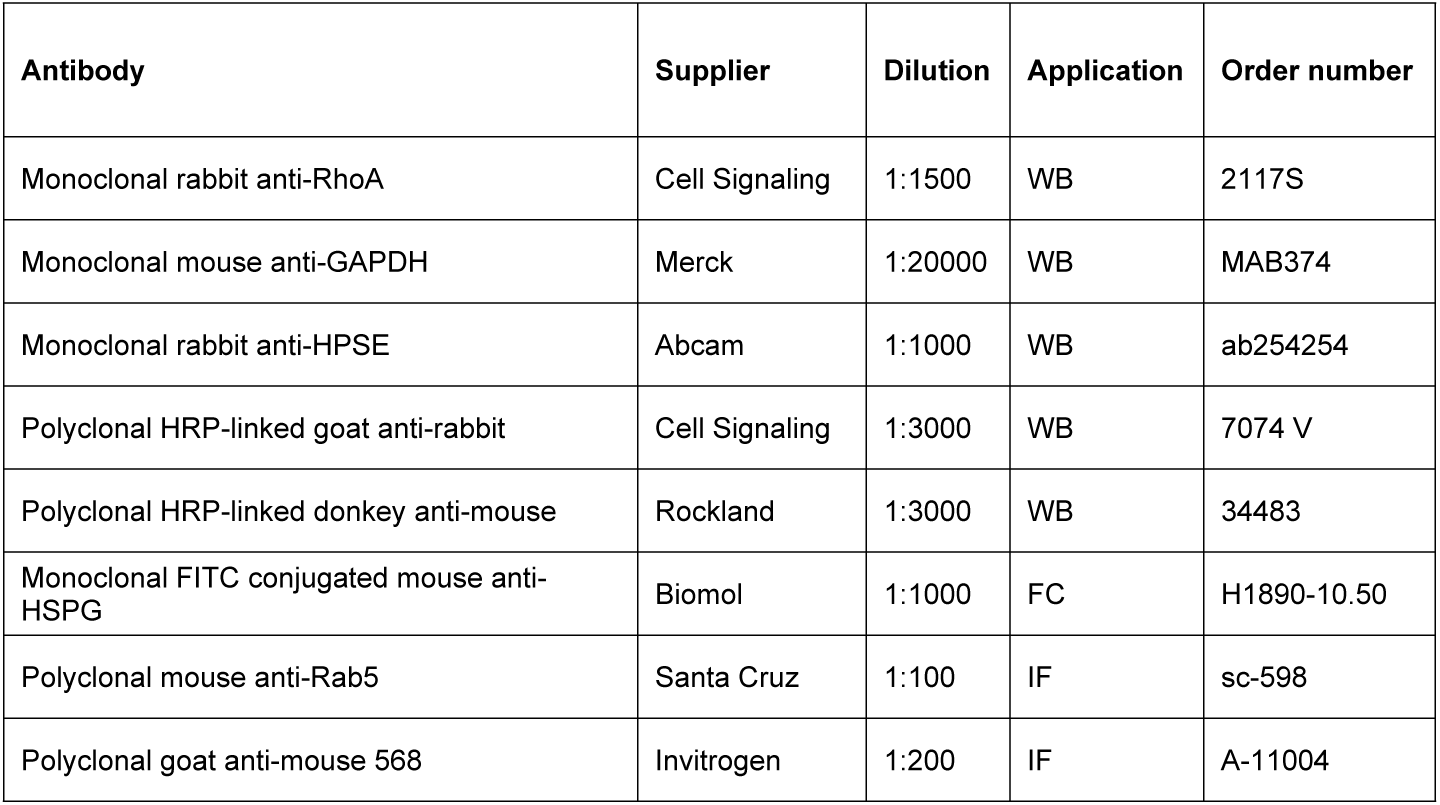

| Antibody | Supplier | Dilution | Application | Order number |
| --- | --- | --- | --- | --- |
| Monoclonal rabbit anti-RhoA | Cell Signaling | 1:1500 | WB | 2117S |
| Monoclonal mouse anti-GAPDH | Merck | 1:20000 | WB | MAB374 |
| Monoclonal rabbit anti-HPSE | Abcam | 1:1000 | WB | ab254254 |
| Polyclonal HRP-linked goat anti-rabbit | Cell Signaling | 1:3000 | WB | 7074 V |
| Polyclonal HRP-linked donkey anti-mouse | Rockland | 1:3000 | WB | 34483 |
| Monoclonal FITC conjugated mouse anti-HSPG | Biomol | 1:1000 | FC | H1890-10.50 |
| Polyclonal mouse anti-Rab5 | Santa Cruz | 1:100 | IF | sc-598 |
| Polyclonal goat anti-mouse 568 | Invitrogen | 1:200 | IF | A-11004 |

List of reagents and their supplier:

| Reagent | Supplier |
| --- | --- |
| Heparin | Sigma (H3149) |
| Fondaparinux | Sigma (SML1240) |
| Dextran sulfate | Sigma (75027) |
| Chondrotin sulfate A | Sigma (C9819) |
| Chondrotin sulfate B (Dermatan sulfate) | Sigma (C3788) |
| Hyaluronic acid | Sigma (53747) |
| DyLight 488 Maleimide-Dye | Thermo Fisher (46602) |
| MES hydrate | Sigma (M8250) |

Mutation sites and primer sequence:

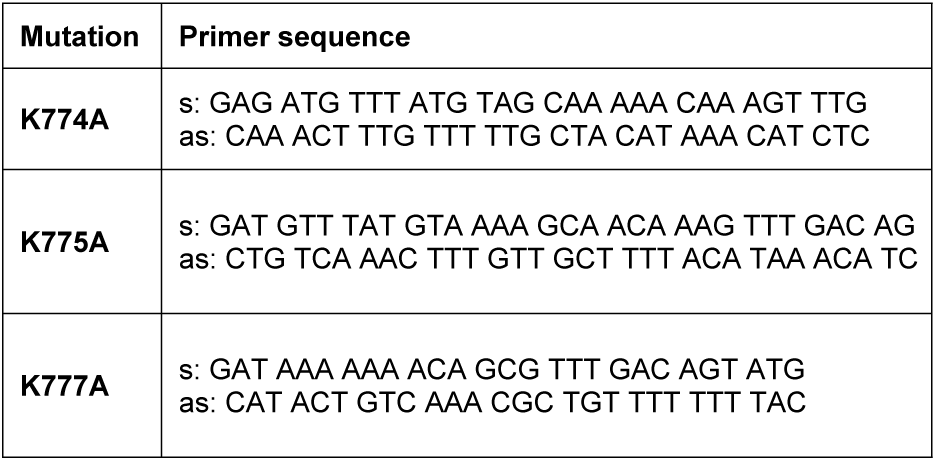

## Methods

### Cell Culture

HeLa (H1, CRL-1958), THP-1, Raw 264.7 and MEF (CF-1) cells were originally obtained from ATCC. Ramos, Molt-4 and J558L were gained from the BIOSS Toolbox. EXT2^-/-^ cells were provided by M. Dong (Harvard Medical School, Boston). Cells were tested negative for mycoplasma contamination monthly. The supplier authenticated cells were obtained from the BIOSS Toolbox. The different cell lines were cultivated in their recommended medium at 37°C with 5% CO_2_ in a humidified atmosphere.

Cell types and needed culture medium.

| Cell type | Culture medium |
| --- | --- |
| **HeLa (H1, CRL-1958)** | DMEM, 10% FBS, 1% NEAA, 4 mM P/S |
| **HeLa EXT2-/-** | DMEM, 10% FBS, 1% NEAA, 4 mM P/S |
| **THP-1** | RPMI 1640, 10% FBS, 4 mM P/S |
| **Raw 264.7** | DMEM, 10% FBS, 4 mM P/S |
| **MEF (CF-1)** | DMEM, 10% FBS, 4 mM P/S |
| **Ramos** | RPMI 1640, 10% FBS, 4 mM P/S |
| **Molt-4** | RPMI 1640, 10% FBS, 4 mM P/S |
| **J558L** | RPMI, 10% FBS, 4 mM P/S |
| **CHO C6** | Ham's F-12, 10% FBS, 4 mM P/S |
| **CHO (pgsA745 (ATCC® CRL2242™))** | Ham's F-12, 10% FBS, 4 mM P/S |

### Cloning, Mutagenesis and Purification of Recombinant CNF Proteins

For generation of GST-CNFY or GST-CNFY_709-1014_ mutants, quick change PCR was performed using specific primers and the plasmid pGEX-2TGL+2 encoding GST-CNFY or GST-CNFY_709-1014_. Plasmids containing the different mutations were transformed into competent *E. coli* TG1. For expression and purification of recombinant GST-CNF proteins, LB-medium containing ampicillin (100 µg/mL) was inoculated with the corresponding bacteria and cultivated overnight at 37°C, 180 rpm. Protein synthesis in the main culture was induced by addition of IPTG (100 µM end concentration) at an optical density (OD_600_) of 0.7. The culture was incubated for additional 20 h at 28°C, 800 rpm. Cells were collected by centrifugation (15 min, 4°C, 6000 × g). The pellet was sonified in lysis buffer (10 mM NaCl, 5 mM MgCl_2_, 5 mM DTT, 1 mM PMSF, 20 mM Tris-HCl, pH 7.3 and 5 µg/mL DNase). Lysates were cleared by centrifugation (45 min, 4°C, 20000 x g) and incubated with Glutathione-Sepharose beads. Protein purification and concentration was analyzed by SDS-PAGE.

### Cloning, Mutagenesis and Purification of Recombinant TcdA protein

The used recombinant TcdA was expressed and purified as described previously (32).

### SiRNA transfection

For silencing the human heparanase (HPSE) mRNA (GenBank ID: 10855; 5’-CTGATGTTGGTCAGCCTCGAA-3’) small interference RNA (siRNA) was used (Qiagen). HeLa-cell were seeded in 6-well plates and cultured to a confluency of 70%. Then culture medium was replaced by 2.5 ml transfection medium (Dulbecco’s Modified Eagle medium + 10% FBS, 1% non-essential amino acids). The siRNA and Lipofectamine™ RNAiMAX Transfection Reagent (Thermo Fisher) were incubated in 500 µl Opti-Mem for 30 min at RT and added to cells for 24 h. The next day, medium was changed to culture medium. In all experiments a negative control siRNAs (5’-AATTCTCCGAACGTGTCACGT-3’) was used. The efficiency of siRNA knockdown was confirmed by Western blotting.

### Cell lysate

Cells were washed with ice-cold PBS and lysed using GST-Fish buffer (10% glycerol, 100 mM NaCl, 1% NP-40, 2 mM MgCl_2_, 1 mM PMSF and 50 mM Tris/HCl, pH 7.4). Lysates were cleared by centrifugation (15 min, 4°C 21000 x g,). Protein amount was determined with the Bradford assay reagent Roti^®^-Quant (Roth).

### SDS Page

Cell lysate was loaded on a 12.5% SDS-Gel. In the case of RhoA deamidation, a 12.5% urea SDS-Gels was used. After protein separation by SDS-PAGE, the gels were either stained with Coomassie or electro transferred onto a PVDF membrane.

### Western Blot

Proteins were transferred onto a PVDF membrane (Carl Roth) using the Wet tank plotting system (Mini Trans-Blot^®^, BioRad). The membrane was blocked with 5% skimmed milk for 1 h at RT. After washing the membrane 3 times with TBS-T for 10 min, primary antibody was added and incubated overnight at 4°C. Unbound antibody was removed by washing 3 times with TBS-T for 10 min. The secondary HRP-linked antibody was subsequently incubated with the membrane for 1 h at RT. Following washing, membranes were developed using SignalFire™ ECL Reagent (Cell signalling) and finally exposed with the imaging system Amersham Imager 600 (GE Healthcare Life Science).

### FACS analysis

Toxins were labelled with DyLight488 Maleimide (Thermo Scientific, Waltham, MA, USA) according to the manufacturer’s instructions. For binding analysis adherent cells were detached from culture plates using 10 mM EDTA in PBS. This was not necessary for suspension cells. After washing, cells were counted and aliquoted for the experiment. For each condition 2 × 10^5^ - 2.5 × 10^5^ cells were used. Cells were incubated for 15-20 min with the indicated DL488-Toxin concentration or 1 h with the FITC-antibody at 4°C. Following washing with ice-cold PBS, cell bound fluorescence was measured using a FACS Melody (BD Bioscience).

### Competition Assay with GAGs or unlabelled CNF

Cells were co-incubated with DL488-GST-CNFY and different concentrations of glycosaminoglycan or unlabelled GST-CNF, respectively as indicated for 20 min at 4°C. Unbound toxin was removed by washing with ice-cold PBS. Cell bound toxin was measured using a FACS Melody (BD Bioscience).

### Confocal images for co-localization

Cells were seeded on coverslips and fixed in 4% para-formaldehyde in PBS, permeabilized with 0,3% Triton X-100, 5% FCS in PBS at RT. Samples were incubated with the Rab5 antibody overnight at 4°C. After washing and incubation with the secondary antibody for 1 h at RT, coverslips were mounted with Prolong diamond + DAPI. Confocal images for co-localization of toxin and the early endosomal marker Rab5 were acquired with a LSM 800 Confocal Laser Scanning Microscope (Zeiss), equipped with a 63x /1.4 NA oil objective and Airyscan detector (Zeiss). Airyscan images were processed by using Zen software (Zeiss)

### Biolayer interferometry (BLI) spectroscopy

The binding affinities between CNF and GAGs were measured using the BLI assay with the BLItz system (ForteBio) as previously described (14). Briefly, 100 μg/mL biotinylated heparin (Sigma-Aldrich, B9806), hyaluronic acid (Sigma, B1557), or cellulose (Creative PEGWorks, CE501) were immobilized onto capture biosensors (Dip and Read Streptavidin, ForteBio). The biosensors were balanced with DPBS (0.5% BSA, w/w) for 30 s, and then exposed to toxins at indicated concentrations for 120 s, followed by dissociation in DPBS (0.5% BSA, w/w) for 180 s. All experiments were performed at RT. Binding affinities (KD) were calculated using the BLItz system software (ForteBio).

### Spin labelling

CNFY was spin labelled with a 40 times excess of MTSL (*S*-(1-oxyl-2,2,5,5-tetramethyl-2,5-dihydro-1H-pyrrol-3-yl)methyl methanesulfonothioate, 100 mM solved in DMSO) overnight at RT. Unbound MTSL was removed by gel filtration on a Superdex 200 10/300 GL column (GE Healthcare). The sample was concentrated using Vivaspin^™^ columns (GE Healthcare) and re-buffered in the measuring buffer (100 mM NaCl, 10% (v/v) Glycerol, 20 mM Tris, pH 7.4/pH 5.0) using a PD-10 column (GE Healthcare). Concentrated sample was loaded into 3.8 mm (outer diameter) synthetic-quartz EPR tubes (Qsil GmbH) and frozen in liquid nitrogen, either directly or after adding heparin.

### PELDOR

Dead-time-free four-pulse electron–electron double resonance (PELDOR) experiments (33) were carried out at Q-band frequency (33.8 GHz) with Bruker Elexsys E580 instrumentation equipped with a PELDOR unit. Microwave pulses were amplified using a 3-W solid-state amplifier. A Bruker dielectric-ring resonator (EN 5107-D2) was used that was immersed in a continuous-flow helium cryostat (CF-935, Oxford Instruments). The temperature during the measurements was 60 K and stabilized to +/-0.1 K by a temperature control unit Oxford Instruments ITC-503. In all measurements, the pump frequency was set to the center of the resonator dip as well as to the maximum of the nitroxide EPR spectrum. The observer frequency was shifted up by approximately 45 MHz. All measurements were performed with observer pulse lengths of 20 and 40 ns for π/2- and π-pulses, respectively, and a pump-pulse length of 40 ns.

PELDOR analyses have been carried out using the analysis tool GloPel (34), which is available free of charge at https://www.radicals.uni-freiburg.de/de/software/glopel-1. Standard Tikhonov regularization was used for the analysis of all PELDOR time traces. All resulting distance distributions have been validated using a method described earlier (35).

### Regression analysis of FACS-derived toxin binding data

Regression analysis was performed with Sigma Plot (Systat Sofware). Toxin binding data from FACS experiments were depicted as scatter plots. Utilizing the Sigma Plot curve fitting tool, “best fit” regression curves were modeled for different binding kinetics. The following regression functions were used for modelling:

One binding site, f(x)=Bmax1*x/(KD1+x).

One binding site / unspecific binding, f(x)=Bmax1*x/(KD1+x)+a*x. Two binding sites, f(x)=Bmax1*x/(KD1+x)+Bmax2*x/(KD2+x).

Two binding sites / unspecific binding: f(x)=Bmax1*x/(KD1+x)+Bmax2*x/(KD2+x)+a*x. Two binding sites / Hill kinetic,

(x)=Bmax1*xh1/(KD1h1+xh1)+Bmax2*xh2/(KD2h2+xh2) | h1 = 1.

Quality of regression curves was evaluated by calculation of residual sum of squares (RSS) values for each curve fit.

## Figures

**Figure S1: Proteoglycan expression of mouse and human cell lines and CNFY activity**.

**(A)** Normalized RNA expression (NX) of different proteoglycans in HeLa, THP-1 and Molt-4 cells. Data were obtained from https://www.proteinatlas.org.

**(B)** Cells were incubated with GST-CNFY and GST-CNFY_C866S_ for 4 h. RhoA was detected by Western blotting using an anti-human RhoA antibody. GAPDH was used as a loading control. The experiments were repeated three times independently with similar results.

**Figure S2: Regression analysis of FACS derived CNFY binding data:** Modelling of a two binding sites Hill kinetic to FACS derived CNFY binding data using the regression function f(x)=B_max1_*x^h1^/(K_D1_^h1^+x^h1^)+B_max2_*x^h2^/(K_D2_^h2^+x^h2^) | h1=1 for (**A)** HeLa cells B_max1_=0.4, K_D1_=11.5, h_1_=1, B_max2_=0.7, K_D2_=57.1, h_2_=2.6 and (**B)** THP-1 cells B_max1_=0.3, K_D1_=19.4, h_1_=1, B_max2_=0.7, K_D2_=78.6, h_2_=3.0. The additive regression function describes binding of the ligand to two receptors and consists of f(x)_1_=B_max1_*x^h1^/(K_D1_^h1^+x^h1^) | h1=1 depicted in blue and f(x)_2_= B_max2_*x^h2^/(K_D2_^h2^+x^h2^) depicted in green.

**Figure S3: Detection of RhoA deamidation by the shift assay:** HeLa cells were incubated with GST-CNFY in the presence of several GAG, respectively as indicated: (dextran sulfate (DxS), heparin (H), fondaparinux (F) and chondroitin sulfate A (CS). RhoA was detected by Western-blotting using an anti-human RhoA antibody. Equal loading was verified by detection of GAPDH. The experiment was repeated three times independently.

**Figure S4: Characterization of the C-terminal heparin-binding motive**.

**(A)** Structure of the CNFY C-terminal domain (709-1014) modeled to the known crystal structure of CNF1 C-terminal domain (709-1014) (36). Amino acids of the heparin-binding motive K774, K775, K777 as well as K786 are highlighted as balls.

**(B)** Flow cytometry analyses of the wildtype and mutant DL488-GST-CNF_709-1014_ (35 nM, 20 min) binding to HeLa cells in the presence or absence of heparin. Values are average of bound fluorescence ± SD normalized to control cells only incubated with DL488-GST-CNFY (n *=* 3). Statistical analysis were performed using one-way ANOVA. ** p < 0.01.

**(C)** Comparative binding curves to immobilized biotin-heparin of CNFY_C866S_, CNFY_709-_ 1014, CNFY_709-1014_ K774A and CNF1_709-1014_. Data are extracted from the experiments shown in Fig. 4.

## Acknowledgments

We thank Jürgen Dumbach, Silke Ludigkeit for excellent technical assistance and SFB 850 (project C2) for funding. SW thanks the SIBW/DFG for financing EPR instrumentation that is operated within the MagRes center of the University of Freiburg.

